# Development of a Novel Method to Fabricate Highly Functional Human Purkinje Networks

**DOI:** 10.1101/2024.10.01.616084

**Authors:** Pengfei Ji, Jeffrey S. Heinle, Ravi K. Birla

**Affiliations:** Laboratory for Regenerative Tissue Repair, Texas Children’s Hospital, Houston, Texas, USA; Center for Congenital Cardiac Research, Texas Children’s Hospital, Houston, Texas, USA; Division of Congenital Heart Surgery, Texas Children’s Hospital, Houston, Texas, USA; Department of Surgery, Baylor College of Medicine, Houston, Texas, USA; Division of Pediatric Surgery, Department of Surgery, Texas Children’s Hospital, Houston, Texas, USA

**Author notes:** **Correspondence:** Ravi K. Birla, PhD, Feigin Center, C.450.07, Texas Children’s Hospital, 1102 Bates Ave., Houston, TX 77030, USA. **E-mail:**.

**Keywords:** Purkinje, heart, electrical activity, bioprinting, cardiomyocytes, co-culture system, tissue engineering, regenerative medicine

## Abstract

**BACKGROUND:** In this study, we present a method to bioengineer functional Purkinje networks using recent advances in laser-based bioprinting.

**METHODS:** A custom bioink as formulated using optimized concentrations of polyethylene glycol diacrylate (PEGDA), gelatin methacryloyl (GELMA), lithium phenyl-2,4,6-trimethylbenzoylphosphinate (LAP), and tartrazine. A standard triangle language (STL) model of Purkinje networks was developed based on the mammalian Purkinje network mapped out using India ink staining. A commercial bioprinter, the Lumen X, from CellInk, was used to bioprint Purkinje networks. The biocompatibility of the bioprinted Purkinje networks was tested using iPSCs from healthy donors. Negative molds of the Purkinje networks were designed to simulate interaction between Purkinje cells and adjacent cardiomyocytes using different degrees of overlap between the two cell types. The negative molds were also shown to be biocompatible, based on the culture of iPSCs derived from healthy donors.

**RESULTS:** We were able to successfully bioprint over 100 Purkinje networks and demonstrate biocompatibility with iPSCs for up to 7 days. Three different configurations of the negative molds were designed and fabricated and all three shown to be biocompatible with iPSCs for up to 7 days. A co-culture system was developed by placing the Purkinje networks in proximity to the negative molds for all three configurations designed.

**CONCLUSION:** Our results demonstrate the ability to bioprint Purkinje networks and molds and provide an in vitro system to study the functional interaction between Purkinje cells and adjacent cardiomyocytes.

## INTRODUCTION

Heart rhythm disorders claim the lives of nearly half a million people in the United States^1^. Abnormal heart rhythms, or arrhythmias, are the result of disturbances in electrophysiology, such as electrical excitation and conduction, leading to decreased mechanical function. Electrical stimulation of the heart is induced by the sinoatrial node of the right atrium, traveling to the atrioventricular node^2^. The atrioventricular node forms an electrical connection between the atria and the ventricles^2^. The atrioventricular node is connected to the Purkinje networks via bundle of His fibers^2^. The Purkinje networks are composed of electrically active cells and fibers that branch from the basal septum to the left and right ventricles^2–4^. These networks are known to send electrical impulses from His-Purkinje cells to cardiomyocytes in the muscles, promoting coordinated contractions of the left and right ventricles^1^. Lack of proper electrical stimulation and conduction can increase or decrease cardiac output negatively, such as tachycardia and bradycardia^5^. Thus, his-Purkinje system is paramount to the heart’s function.

Common therapies and treatments for cardiac arrythmias caused by the His-Purkinje system (HPS) include cardioverter-defibrillators, antiarrhythmic drugs, and catheter ablation, and pacemakers that attach to the His bundle. Pharmaceutical therapies include beta blockers and calcium channel antagonists but are restricted to patients without other heart complications. Despite amiodarone drug’s strong effectiveness in the maintenance of heart rhythm, with an 11% recurrence of fibrillation in patients taking the drug, its side effects increase with higher doses; thus, long term use is not recommended. Catheter ablation is a viable option for patients with less server cases of congenital heart disease, but those with more severe complications have a higher recurrence rate. Most treatments involve dampening the symptoms and effects, but unable to stop reoccurrence of arrhythmias. There are still clear gaps in creating a standardized treatment strategy due to the limited knowledge of Purkinje networks.

Purkinje networks are complex structures that are limited in their ability to be imaged thus difficult to reconstruct *in vitro*^6^. However, studies on Purkinje network remodeling have utilized animal models, where the Purkinje networks were modified *in vivo*. For example, congestive heart failure was promoted in rabbits via ventricle and pressure overload. This induced impaired function of the Purkinje Network and remodeling of the network itself, concluding that the remodeling of PN caused increased arrhythmic events (Chiu, 2008 #1). However, it has been noted that networks differ anatomically among different mammalian species, thus animal models aren’t accurate in understanding the human Purkinje network (PN). Previous investigators have constructed Purkinje networks that only mimicked the characteristics of the network rather than the anatomical structure. Thus, a more functional and accurate model for PN is needed.

Much of the field of cardiac tissue engineering has focused on developing methods to fabricate components of the heart, including heart muscle tissue^7–9^, vascular grafts^10^, biological pumps^11^, ventricles^12–16^ and whole hearts^17^. In these studies, the goal has always been to replicate the contractility of the heart. This is of paramount importance. However, there have been very few studies focused on developing methods to recreate the complex geometry of the mammalian Purkinje network. This is not due to the lack of significance of such work, but rather a reflection on the complexity of the mammalian Purkinje networks and the lack of fabrication methods available to replicate such a complex geometry.

The ability of bioengineer Purkinje networks in the lab has several advantages and such a system can be used for many different applications^18^. First, bioengineered three-dimensional (3D) Purkinje networks can be used to study the electrical conduction patterns in normal and diseased hearts; with the recent advent of method to reprogram human induced pluripotent stem cells (iPSCs) to Purkinje cells, this provides a very powerful tool to study patient specific electrical activity. With the absence of Purkinje networks, such studies can only be conducted in two-dimensional (2D) monolayer systems, which lack the directionality of the Purkinje networks. Furthermore, the interaction of Purkinje networks with adjacent cardiomyocytes is an important determinant of heart function. The electrical activity of Purkinje networks in the form of depolarization waves interact with cardiomyocytes in the well-studied excitation-coupling (E-C) coupling. However, there are currently no *in vitro* models of such a system, due to the lack of suitable 3D Purkinje systems. Such a system will be powerful to investigate the effect of specific genetic defects on the electrical conduction in isolated *in vitro* Purkinje networks and subsequently, the impact on cardiomyocytes.

In this study, we sought out to develop technology to fabricate mammalian Purkinje networks, taking advantage of recent advances in high resolution laser based bioprinting. Our object was to develop and optimize custom bioink formulations, optimize bioprinting parameters, develop technology to bioprint mammalian Purkinje networks and develop a co-culture system for Purkinje networks with cardiomyocytes.

## Materials and Methods

All chemicals, reagents, and solvents were purchased from Sigma-Aldrich (St. Louis, MO, USA) unless otherwise specified.

### Generating STL Files for Purkinje Networks

A recent publication by Dr. Vilhena very elegantly described the bovine Purkinje network of the left ventricle^6^. India ink was used to visualize the Purkinje network and furthermore, the staining protocol was optimized to provide a detailed image of the Purkinje network (**Figure 1A**); in this model, the left bundle branch feeds into the left ventricle and forms a very complex branching network of Purkinje fibers which allows synchronized contractions of the heart. Using the anatomical structure in this manuscript, a 3D model of the Purkinje network (**Figure 1B**) was created using SolidWorks Computer Aided Design ^19^ software.

**Figure 1:**
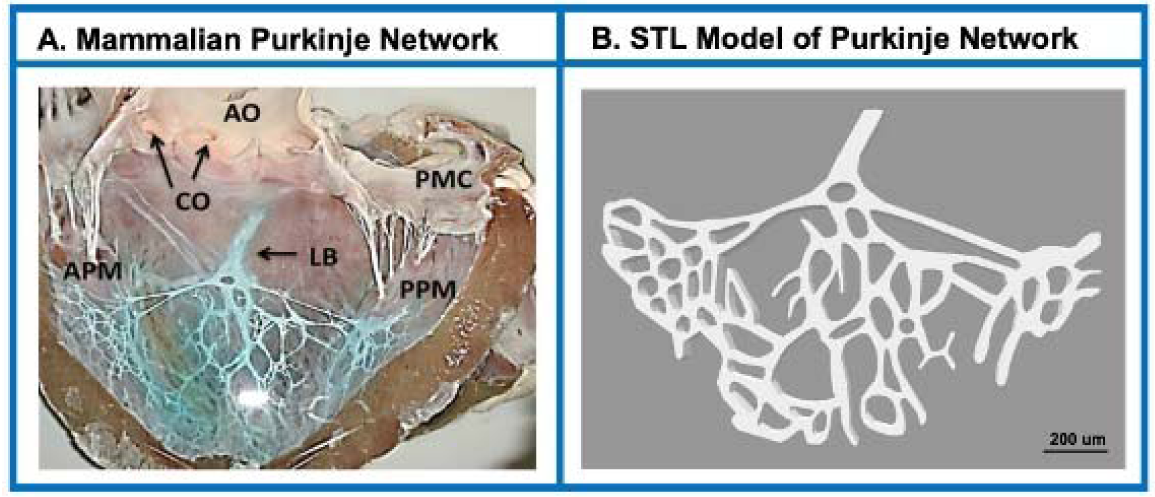
Development of STL models for Purkinje networks: CAD generated STL file of Purkinje networks based on the bovine model.

### Bioink Formulation

We used a custom blend of methacrylated gelma (Advanced Biomatrix, Carlsbad, CA, USA), dissolved in PBS at a stock concentration of 20% and stored at -20°C and used at a final concentration of 5% in the bioink formulation. Polyethylene glycol diacrylate, molecular weight of 3,400 (Alfa Aesar, Tewksbury, MA, USA) dissolved in PBS at a stock concentration of 50% and stored at -20°C and used at a final concentration of 10% in the bioink formulation. A stock solution of lithium phenyl-2,4,6-trimethylbenzoylphosphinate (LAP), (Sigma) was made at a concentration of 1% in PBS and stored at -20°C and used at a concentration of 200 mM in the bioink solution. A stock solution of tartrazine (Sigma) was made up in PBS at a concentration of 10%, stored at room temperature and used at a concentration of 100 mM in the bioink formulation. 10 ml of bioink was prepared freshly prior to every print and used immediately.

### Bioprinting Parameters for Material Based Purkinje Networks

We used a commercially available laser based bioprinter for our applications, the Lumen X from CellInk. The heating element on the printer stage was activated and set at 60°C. Once the stage reached the operating temperature, the bioink was transferred to the stage and allowed to equilibrate at the operating temperature for 15 minutes. Printing was conducted using a laser intensity of 60% and exposure time of 30 second. Individual prints were washed with PBS and maintained in cell culture media in a 37°C cell culture incubator.

### Culture and Expansion of iPSC’s

A control line of iPSC cells was purchased from the BCM stem cell core and maintained on Matrigel coated tissue culture plates using mTeSR^TM^ Plus (Stem Cell Technologies, Vancouver, BC, Canada) with media changes every day.

### Bioprinting Purkinje Networks with iPSCs

iPSCs were mixed with our bioink formulation at a density of 1 million cells per ml and used to bioprint Purkinje networks using at the same parameters described for material-based printing, though the temperature of the printing stage was maintained at room temperature.

### Development of co-culture system

Solidworks was used to generate a negative of the Purkinje networks and varying degrees of overlap were introduced by removing specific geometric objects from within the design. A total of three models were generated. First, with no changes to the inner structures and represents a true negative of the Purkinje networks. Second, 50% of the inner structure removed from the design and represents and intermediate design. Third, the entire geometric elements removed from the inner of the design, representing contact at the boundaries.

### Fixing and Staining of Cellularized Purkinje Networks

Purkinje networks with iPSCs were first transferred onto 6-well plate and then fixed using 4% paraformaldehyde for 24 hours. Next, the 4% paraformaldehyde was removed, and the Purkinje networks washed three times with PBS. The tissue culture plate was loaded with 1 ml/well blocking buffer 3%BSA/0.1% Triton-X in PBS (ThermoFisher, Waltham, MA, USA) and incubated for 2 hours at room temperature. After that, the blocking buffer was removed, and cocktails were made by DAPI solution 1:5000 in PBS. VECTASHIELD® PLUS Antifade Mounting Medium with DAPI (Vector Laboratories, Newark, CA, USA) and SSEA- 4- Alexa Fluor 488, 1:20 BD Pharmingen™ Alexa Fluor® 488 Mouse anti-SSEA-4 Clone MC813-70 (RUO) (BD Biosciences, Franklin Lakes, NJ, USA) 2 ml/well solution. Staining was for 8 hours. Next, the Purkinje networks were washed three times, and then transferred to a specialized 6-well glass bottom plate for imaging (Cellvis, Mountain View, CA, USA). The Purkinje networks in were imaged and processed by CellVoyager CV8000 Yokogawa confocal scanner microscope. The z-depth are selected at -150um and 350um, and the thickness of the scanner layer is 6.5um. The lasers of wavelength are 405nm for DAPI and 488nm for the SSEA-4- Alexa Fluor 488, and the objective is chosen the 4x lens to observe the Purkinje network. The images were conducted, merged and analyzed by CellPathfinder (Yokogawa, high-content analysis software) to show the 2/3 D aligned images.

## Results

### Fabrication of Purkinje Networks

The method to generate Purkinje networks, as shown in **Figure 2**, was reproducible and till date we have been able to reproducibly generate over 100 Purkinje networks. The success rate of generating Purkinje networks was 100%, as determined by visual inspection of the bioprinted structure, showcasing the ability to maintain an intact 3D structure. In addition, once hydrated in PBS, the bioprinted Purkinje networks maintained a stable geometry when maintained in a cell culture incubator for up to 7 days (longer time points were not tested).

**Figure 2:**
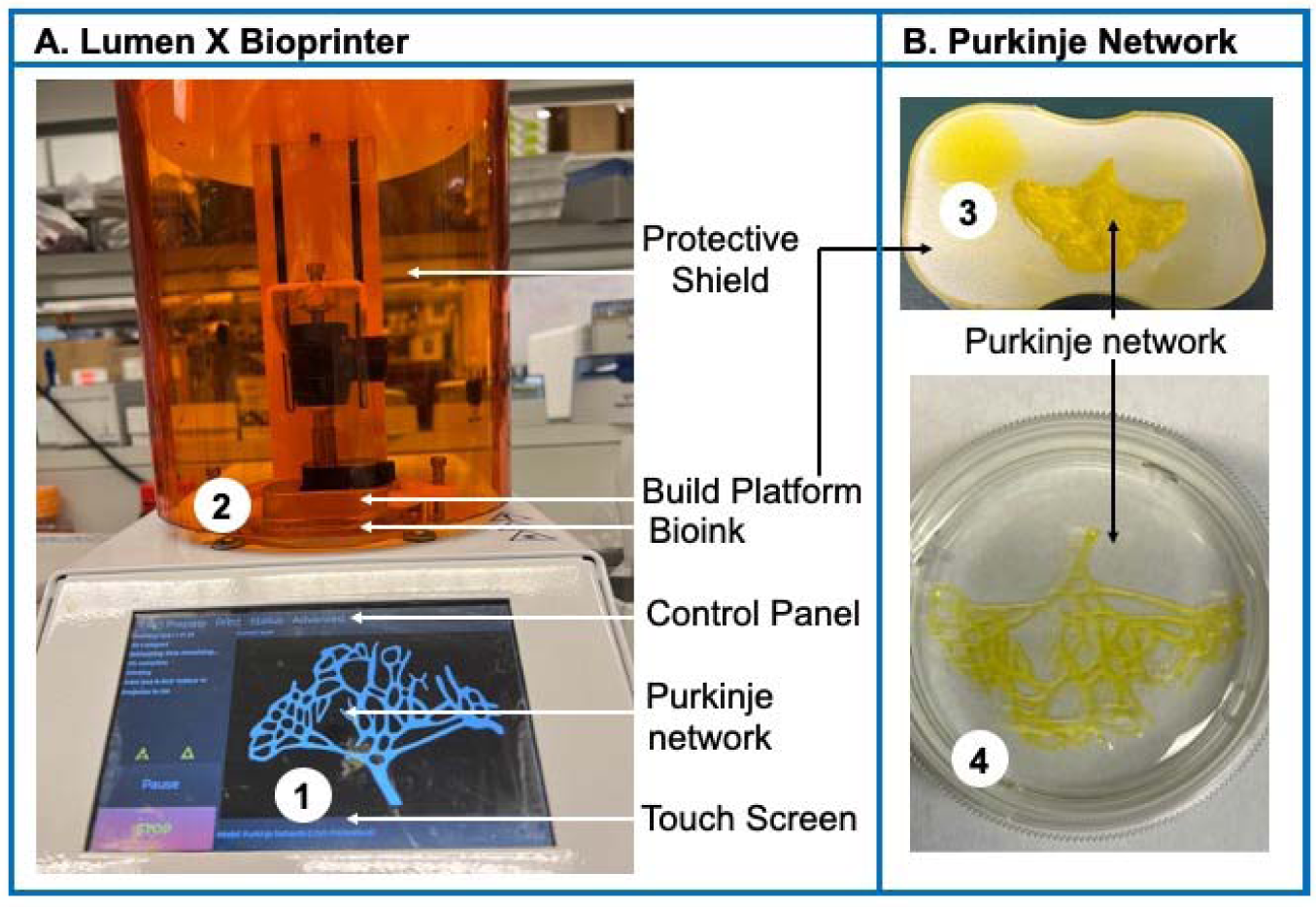
Workflow for printing Purkinje networks: **(A) Lumen X Bioprinter:** A commercially available was used, the Lumen X from CellInk. Major components of the bioprinter are labeled on the image. **(B) Purkinje Network:** The Purkinje network remains adhered to the printing block and then separated and transferred to a tissue culture plate.

### Generation of Cellularized Purkinje Networks

As shown in **Figure 3**, the Purkinje networks supported the culture of live iPSC for up to 4 days, as assessed using staining with DAPI. Confocal imaging was used to visualize the distribution of live cells within the Purkinje networks and iPSCs were noted throughout the thickness of the 3D structure and uniformly distributed within each layer. These results serve to demonstrate the compatibility of bioengineered Purkinje networks with cells and ability to support cell survival and viability.

**Figure 3:**
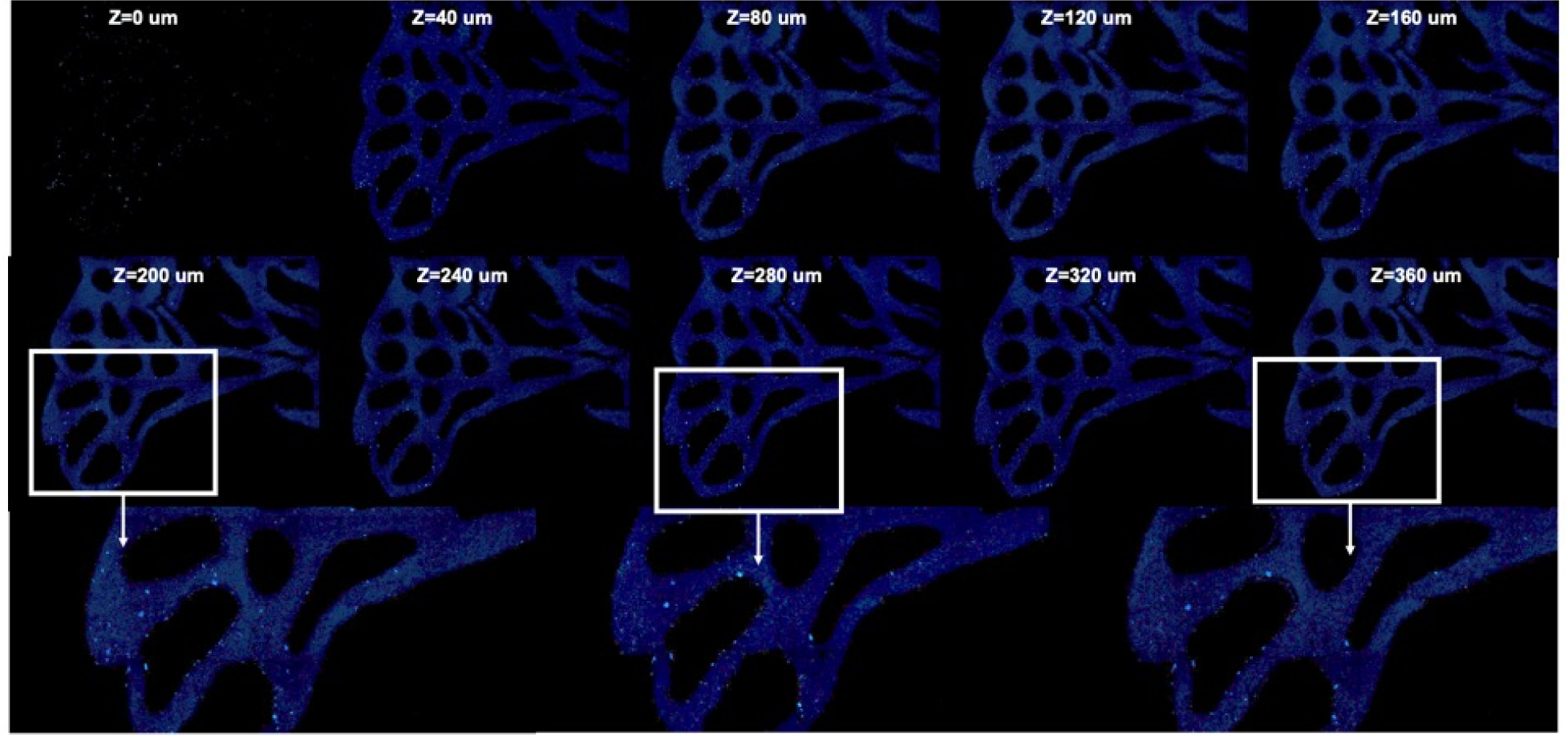
Supporting iPSC culture in Purkinje networks: Serial sections using a confocal microscope was used to visual the distribution of DAPI stained iPSC cells.

### Development of Co-culture systems with Purkinje Networks

To simulate the interaction of Purkinje cells with adjacent cardiomyocytes, we developed a mold that represents the negative of the Purkinje networks. Three variations of the cardiomyocyte molds were generated, representing differential degrees of overlap between the two cell types, **Figure 4**. In each of the three cases, a coloring scheme was used to visualize the Purkinje networks against the cardiomyocyte molds and in all three cases, there was a perfect fit between the two molds. Furthermore, the cardiomyocyte mold was shown to support iPSC culture, **Figure 5**, though this was expected as the bioink was the same one used to generate Purkinje networks. Dual labeling of the iPSCs within the Purkinje and cardiomyocyte molds was used validate the co-culture system, **Figure 6**.

**Figure 4:**
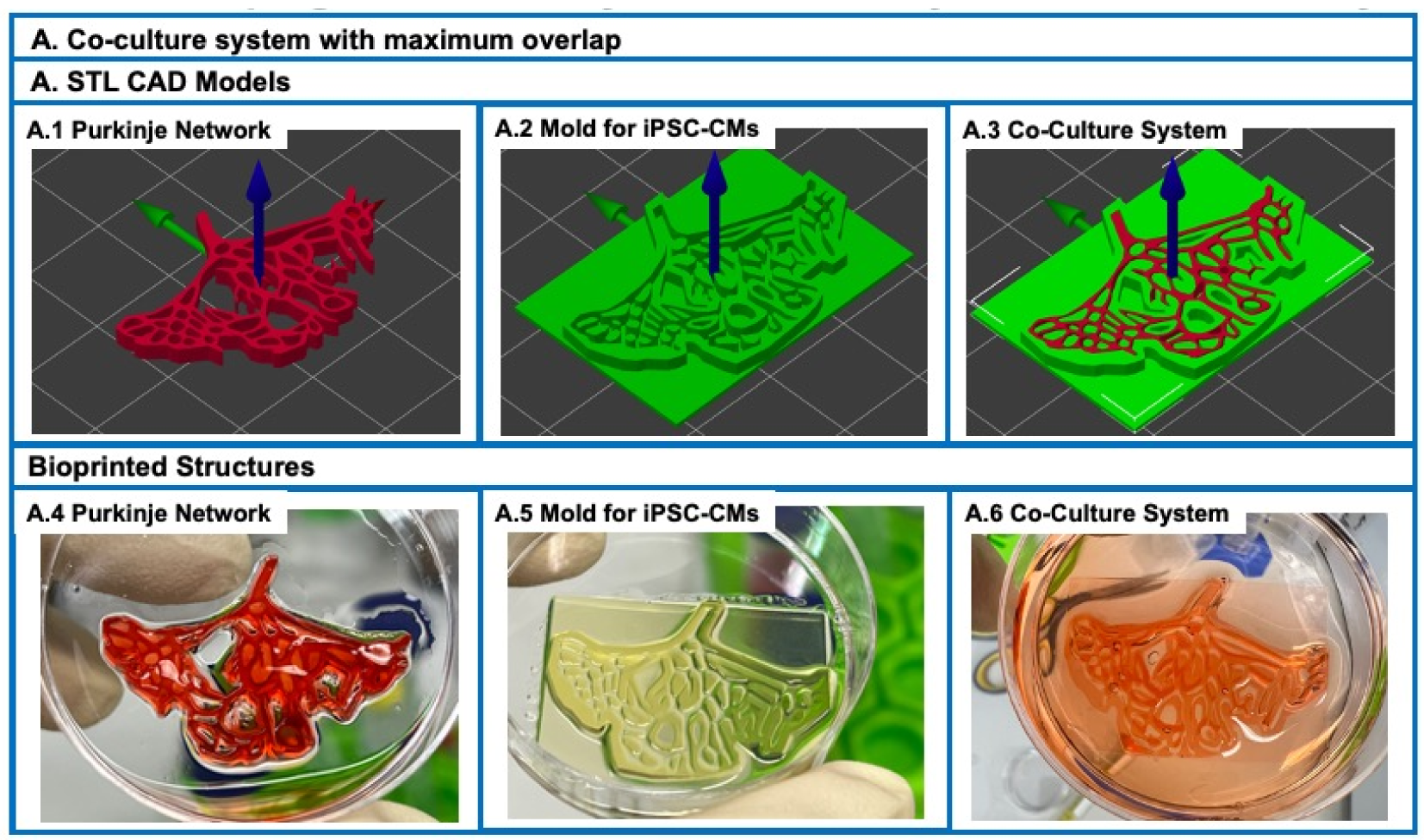
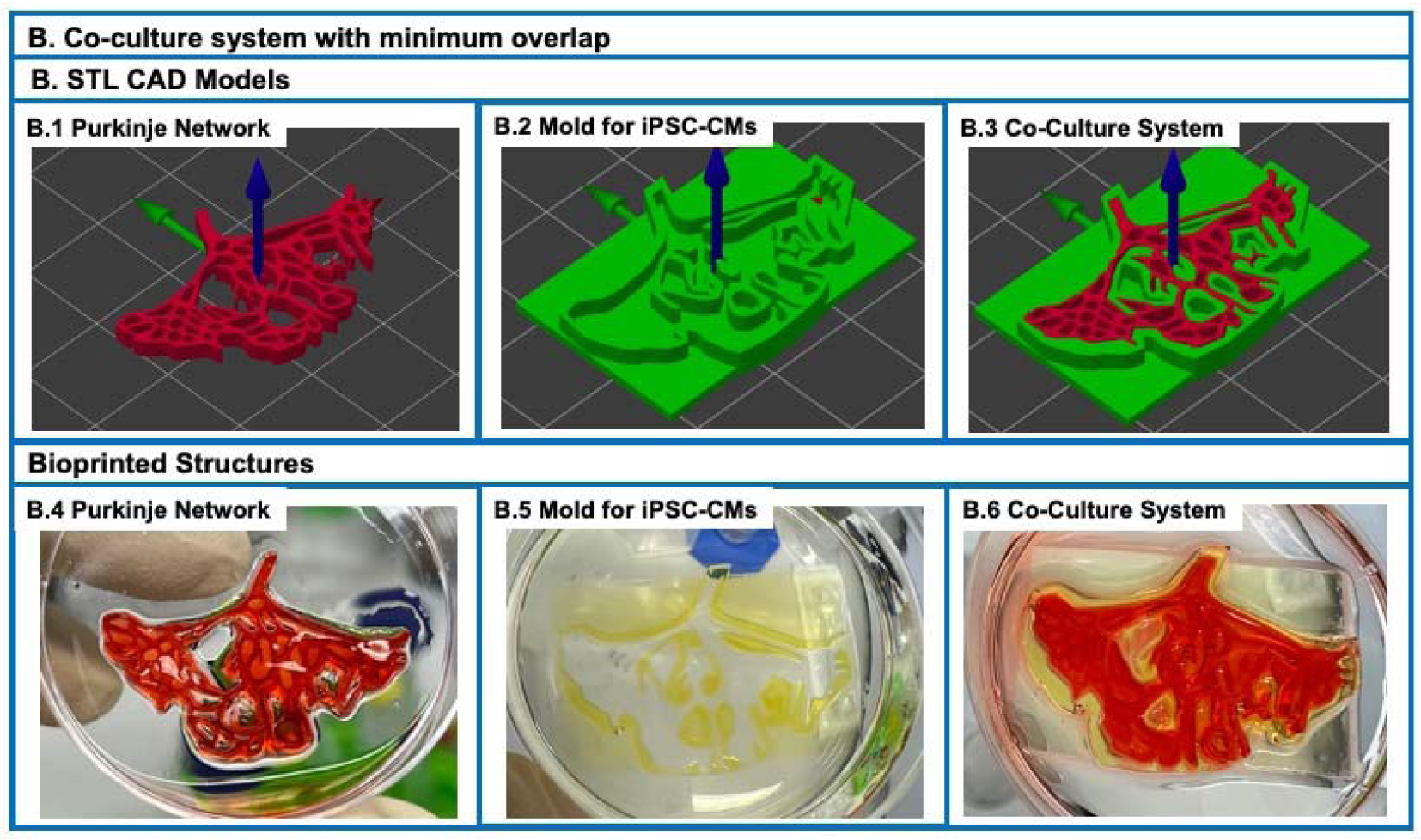
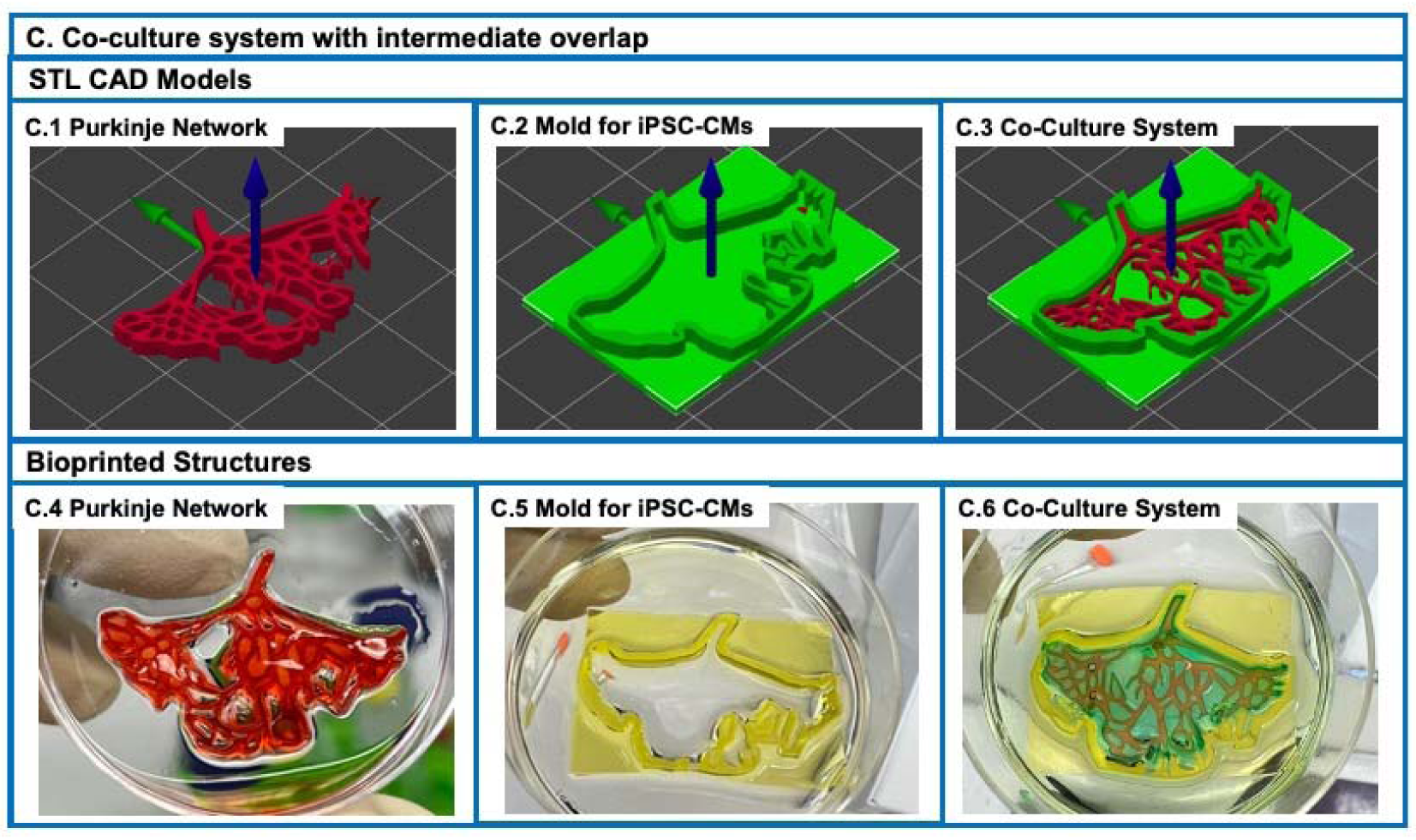
Co-culture system for Purkinje networks: **(A) Co-culture System with maximum overlap:** The co-culture system was designed as an exact negative of the Purkinje network to maximum the degree of overlap between the Purkinje network and the negative mold. **(B) Co-culture system with intermediate overlap:** A fraction of the internal structure of the mold was retained to provide intermediate overlap between the Purkinje network and mold. **(C) Co-culture system with minimum overlap:** The inner content of the negative mold was removed to provide the minimum overlap between the Purkinje network and the negative mold. For all three cases (maximum, intermediate and minimum overlap) the images in the top panel showcase the STL files of the Purkinje network, the negative mold and both the Purkinje network and the negative together. For visualization of all three configurations (maximum, intermediate and minimum overlap) the molds were labeled with dyes and then the two molds placed together to validate the co-culture system.

**Figure 5:**
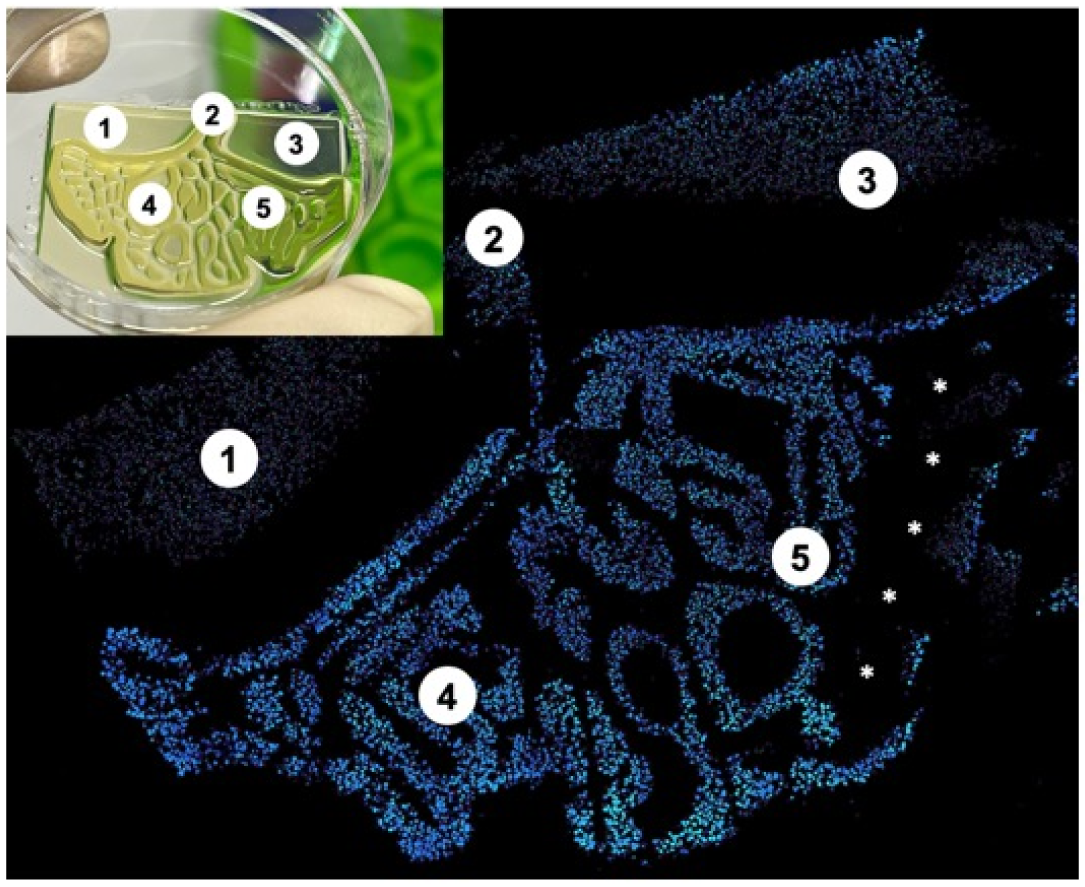
Culturing iPSCs within co-culture molds: Serial sections using a confocal microscope was used to visual the distribution of DAPI stained iPSC cells. The insert on the top left panel shows the bioprinted structure. Five points, labelled as 1, 2, 3, 4, 5 have used to identify different regions on the bioprinted Purkinje network and the labelled images.

**Figure 6:**
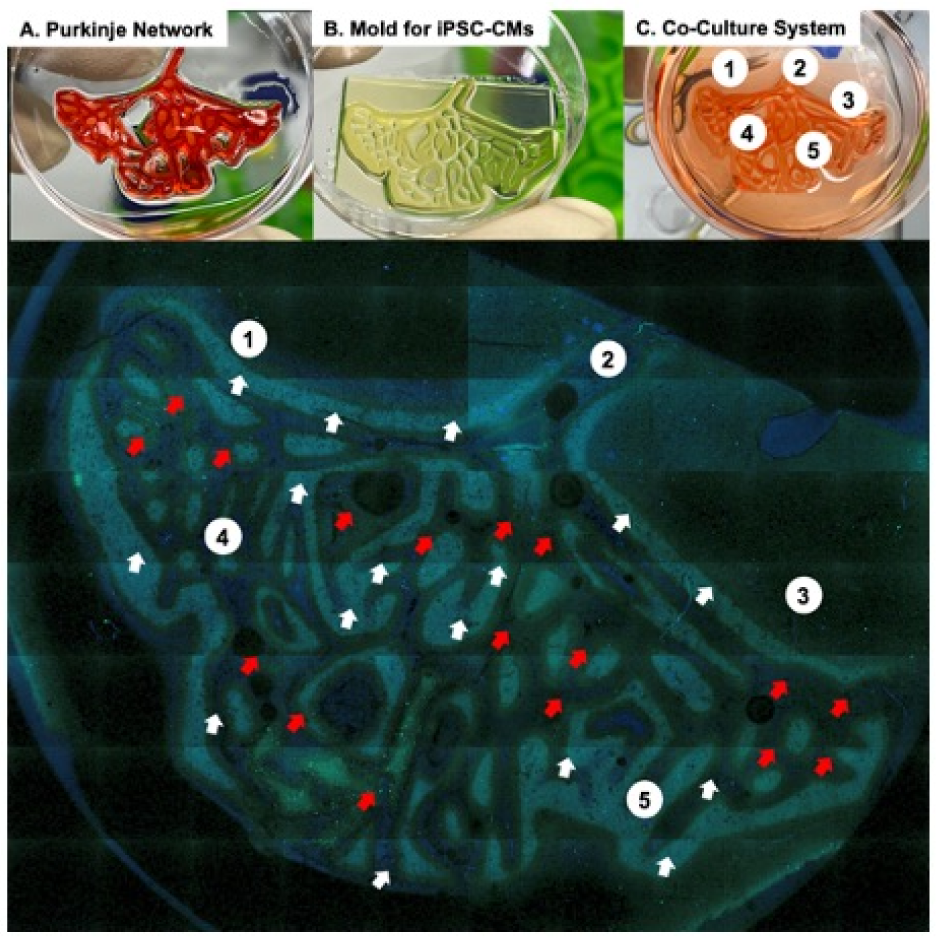
Dual Labeling for Co-culture system: **(A) Purkinje Network:** Bioprinted Purkinje network. **(B) Mold for iPSC-CMs:** The mold with the maximum overlap between the Purkinje networks and the mold was selected. The mold is designed is to simulate the iPSC-CMs. **(C) Co-culture system:** The bioprinted Purkinje network and the mold were placed together. The lower images show the results of the imaging of the co-culture system. iPSCs labeled by DAPI or SSEA-4. Mold with iPSCs were stained by DAPI, and Purkinje Network was stained by SSEA-4-AF488. Representative scanner image showing the merged image of the mold and Purkinje Network together. iPSCs in Purkinje network was SSEA-4-AF488(green), and iPSCs in mold were nuclear counterstain (DAPI, blue). The images were obtained using a 4x objective.

## Discussion

This is a technology development study designed to bioengineer Purkinje networks. Much of the research in the field of cardiac tissue engineering has focused on replacing the contractility of the heart. This work has resulted in the development of heart muscle tissue, biological pumps, ventricles, and whole hearts. While this work has defined the field, there has been a notable absence in technology to recreate the cardiac conduction system, namely, the Purkinje networks within the heart. The Purkinje networks are a critical determinant of heart muscle function and interact with heart muscle tissue to determine the functional performance of the mammalian heart. While the Purkinje networks represent an important component of the mammalian heart, efforts to bioengineer Purkinje networks have been limited. This is likely due to the inherent complexity of the Purkinje networks, with its complex branching pattern and the lack of suitable fabricate methods for this specific application. In this study, we sought to develop a method to bioengineer mammalian Purkinje networks and develop a co-culture system to study the functional interaction between Purkinje networks and cardiomyocytes.

To support bioprinting Purkinje networks, we made use of recent advances in laser based bioprinting and selected this technology for its increased resolution, relative to extrusion-based methods. We selected GELMA and PEGDA as our based materials for our bioink and optimized the concentration of these two components; PEGDA provides mechanical strength while GELMA provides biological activity. LAP was used our cross-linking agent and tartrazine for selective blocking of the light source; bioprinting parameters were optimized to support the fabrication of Purkinje networks. Though time consuming, this optimization process was required to determine optimal biomaterial and bioprinting parameters to generate Purkinje networks. This optimization process was successful and resulted defining optimal parameters and subsequently, over one hundred Purkinje networks have been printed with a 100% success rate.

We confirmed the ability of the bioengineered Purkinje networks to support the culture of cells, using control undifferentiated iPSCs as our test bed. While the Purkinje networks are designed to support the culture of his-Purkinje cells, our lab currently does not have access to these cells. Therefore, as an alternative, undifferentiated iPSCs were considered a viable option. Purkinje networks were shown to support the culture of iPSCs for a period of up to 4 days, assessed using confocal microscopy of DAPI stained cells. These results served to demonstrate the utility of bioengineered Purkinje networks to support the culture of cells and provide validation of these use of this model to bioengineer functional Purkinje networks populated with his-Purkinje cells.

Next, our goal was to develop a co-culture system to investigate the functional interaction of his-Purkinje cells with cardiomyocytes. While it is known that Purkinje cells influence the behavior of cardiomyocytes, the mechanisms by which this occurs remains largely unknown, primarily due to the lack of *in vitro* models to study this. In our work, we developed three variations of the co-culture system, with varying degrees of overlap, all of which were shown to be compatible with the Purkinje networks. As was the case for Purkinje networks, our negative molds were also populated with undifferentiated iPSCs to demonstrate cell culture and were shown to be successful. While populating the negative molds with iPSC derived cardiomyocytes would have been more suitable, the use of undifferentiated iPSCs served to demonstrate compatibility with cells.

The current study demonstrated our ability to bioengineer Purkinje networks and our ability to develop co-culture systems for Purkinje cells with cardiomyocytes. Further development of this technology is needed to address several gaps in the current study. The Purkinje networks need to be populated with iPSC derived his-Purkinje cells and the negative molds need to be populated with iPSC derived cardiomyocytes. Bioreactors need to be fabricated to support electromechanical stimulation of the Purkinje networks to support the maturation of his-Purkinje cells and subsequent maturation of the bioengineered Purkinje networks. While there are additional steps that need to be completed in order to support the development of this technology, the current study provides the platform for this additional work.

There are numerous applications of this technology. Bioengineered Purkinje networks can be used as an in vitro model system to study the directional conduction of electrical waves in normal physiology and pathology of the cardiac conduction system. These bioengineered Purkinje networks can be combined with contractile 3D heart muscle patches to provide both contractile and electrically active heart muscle tissue. Bioengineered Purkinje networks can also be used as a building block for bioengineered hearts.

There are some limitations associated with this study. While significant progress has been demonstrated in the bioprinting of Purkinje networks and subsequent co-culture systems with cardiomyocytes, these systems have not been tested with the correct cell types. As initial proof of concept, both the Purkinje networks and the negative molds for the cardiomyocytes were tested with undifferentiated iPSCs; this work demonstrated the compatibility of bioengineered structures with iPSCs. However, the more pertinent cell types will be the functional cell types. These include iPSC derived his-Purkinje cells and cardiomyocytes and will be the focus of further studies.

This study describes a new and novel method to bioengineer Purkinje networks and subsequent co-culture with cardiomyocytes. Many of the concepts are first in the field of cardiac tissue engineering, similar to where the field started over two decades ago. It is without doubt that in the near future, Purkinje bioengineering will expand to become a dominant component of cardiac tissue engineering and perhaps, transform into a field by itself. The current study will serve as the seminal platform for the development of the field of Purkinje bioengineering and prove to be instrumental in developing and expanding this field.

## Abbreviations

HPS: His-Purkinje system
3D: three-dimensional
2D: two-dimensional
CAD: Computer Aided Design
LAP: lithium phenyl-2,4,6-trimethylbenzoylphosphinate
GELMA: gelatin methacryloyl
PEGDA: polyethylene glycol diacrylate
STL: standard triangle language
PN: Purkinje Network
iPSCs: Induced pluripotent stem cells
E-C: excitation-contraction

## Acknowledgments

RKB would like to acknowledge financial support from the Division of Congenital Heart Surgery, at Texas Children’s Hospital.

## Funding and/or Conflicts of interests/Competing interests

The authors do not have any conflicts of interest or competing interests to declare.

